# Microstate in rat’s EEG: a proof of concept study

**DOI:** 10.1101/2025.06.17.660121

**Authors:** Vaclava Piorecka, Cestmir Vejmola, Petra Peskova, Marek Piorecky, Stanislav Jiricek, Vlastimil Koudelka, Inga Griskova-Bulanova, Tomas Palenicek

## Abstract

The electroencephalogram (EEG) reflecting brain activity can be characterized through brief periods of stable neural activity patterns that recur over time and are referred to as microstates. Microstates are related to a range of cognitive processes, and their analysis has become an increasingly popular tool for studying human brain function. While microstates have been extensively studied in humans, their presence and characteristics in animal models have yet to be as thoroughly investigated. This study aims to address this gap by detecting and characterizing microstates in EEGs of rats collected using a superficial electrode system corresponding to homological areas of the human 10-20 system. Specifically, we demonstrate the presence of microstates in rats’ EEG; those can be captured by the same metrics as in humans. We define these microstates, describe them through topology and parameters, and identify the EEG frequency bands and intracranial sources that predominantly determine microstate topography. These findings have important implications for the use of microstates as a preclinical tool for investigating brain functions, detecting new biomarkers of brain diseases, and translating this knowledge to humans.

## 1. Introduction

Electroencephalography (EEG) is a non-invasive technique that allows the recording of brain activity with high temporal resolution. In the past few decades, EEG microstate analysis has become a popular tool for investigating the functional organization of the human brain [1, 2].

Microstates are brief periods of stable topographical configurations of scalp voltage maps lasting a few hundred milliseconds (40–120 ms). Typically, around or more than 70 % of the variance in the human EEG signal during resting state (no active task) is attributed to four to seven prototypical microstate types, denoted as A, B, C, D, and E, F, G [3]. Each microstate class is characterized by distinct topographic appearances [3–5] and associated functional correlates [3]. It is assumed that microstates represent the brain’s functional states, during which various brain regions communicate in a coordinated manner [6]. Indeed, several studies have shown that intracortical sources of various microstates spatially correspond to the hubs of fundamental resting state networks [7, 8]. Thus, the temporal features of the microstates, such as their average duration in milliseconds, frequency of occurrences per second, and their percentage contribution to the EEG signal, can serve as sensitive indicators of various momentary mental states and stable trait characteristics [9–12]. Additionally, these temporal characteristics could serve as potential biomarkers for mental and neurological disorders, making the approach a valid tool for pre-clinical and clinical research.

Given this emerging significance of microstates, it is striking that they have so far been overlooked in studies on animal models, which would not only allow an in-depth understanding of microstates through feasible manipulations (pharmacological, genetic, behavioral, or neural manipulations such as optogenetics or deep-brain electrode stimulation) but also stand as a valuable tool in translational research. To the best of our knowledge, only four up-to-date studies reported the analysis of microstates in rodent models. Mégevand et al. observed that the somatosensory-evoked responses in mice can be characterized by a series of distinct cortical map configurations, each one remaining stable for a given period of time and then quickly changing into a new configuration in which it remained stable again [13]. Two other studies focused on the identification of microstates for the estimation of the spatiotemporal properties of neuronal ensembles in LFP recordings. Authors reported the existence of a kind of “mesoscale” microstates that exhibit high spatial coherence, temporal stability, and frequency-specific modulation. These microstates were modulated by the behavioral state, such as movement or exploration [14], and by depth of anesthesia [15]. Most recently, Boyce et al. [16] studied EEG microstate dynamics during resting wakefulness in mice from the multi-EEG surface recordings and demonstrated that local cortical neural assembly activity coordinates with global brain dynamics during wakefulness but not during sleep.

Although the above studies suggested that microstates are present in rodents like humans, none used a homologous electrode system and rigorously described microstate parameters using appropriate metrics to make the method translationally usable. Here, we analyzed signals from 30 rats recorded using 21 electrodes placed in regions homologous to the human 10-20 EEG system and adopted analysis approaches used in human research to achieve. We employed several criteria to determine the optimal number of microstates, and, following Mishra et al. [14], utilized surrogate data analysis to contrast the global explained variance (GEV) parameter between randomly shuffled data and actual EEG recordings. We described the characteristics of the optimal microstates, including their topographies, duration, occurrence, and GEV values. We also explored their correlation structure in spatial and temporal domains. Additionally, following human EEG microstate studies [17, 18], we aimed to determine the brain areas involved in the generation of microstates. This knowledge is crucial for the translational utility of the microstate approach from humans to animal models and for understanding the neural basis of brain function across species.

## 2. Materials and Methods

### 2.1. Data collection

All experiments were carried out on adult male Wistar rats (SPF animals; Velaz, Czech Republic) weighing 280–300 g. 30 animals in total were used. Each animal was tested only once. Access to water and a standardized diet was ad libitum. Ethical approval was given by the National Committee for the Care and Use of Laboratory Animals, CZ, and European Union guidelines and principles (86/609/EU) were adhered to.

Rats were stereotactically implanted with 21 gold-plated electrodes (Mill-Max) under general isoflurane anesthesia (2.5% concentration). Electrodes were implanted onto the surface of the cortex in homologous frontal, parietal, and temporal regions of the right and left hemispheres. The coordinates were designed to homologically and functionally match the human 10-20 electrode system. Coordinates were taken from the Paxinos rat brain atlas [19]. Their positions are listed in Figure 1. The reference electrode was implanted above the olfactory bulb, and the ground electrode was placed subcutaneously in the occipital region. All electrodes were fixed to the skull with dental cement. The whole procedure took 45 min on average. Postoperatively, the rats were treated with ketoprofen (5 mg/kg, s.c.) and housed individually (to prevent biting of the implant) and handled daily to habituate them to the manipulations. One day before recording, a connector was mounted to the electrodes under short-term anesthesia and fixed with dental cement.

**Fig. 1.**
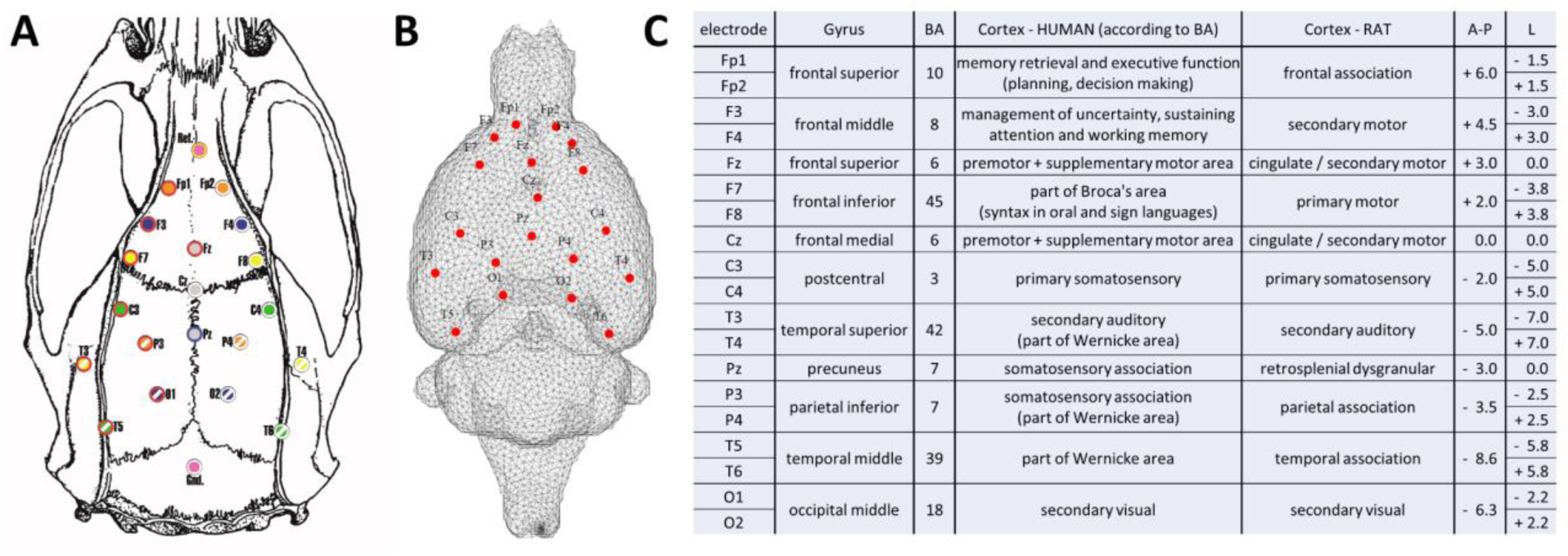
Electrode layout shown on the rat skull **(A)** and brain mesh model **(B)** used for forward modeling. The table **(C)** shows the electrodes with their coordinates and locations in the original 10-20 system distribution in humans, with homologous regions of selected coordinates for the rat system. BA, Brodmann areas; A-P, antero-posterior; L, lateral. Coordinates are given in millimeters, according to [19].

The experiments were conducted during the daytime, between 7:30 and 13:00, 7 days after surgery. Approximately 15 minutes before the experiment began, the animals were connected to the EEG system in their home cages. Subsequently, a 100-minute recording was taken while the rats were allowed to move freely around the cage. Raw EEG signals were acquired using a BioSDA09 standard 32-channel digital EEG amplifier (M&I Ltd., Prague, Czech Republic) at a sampling rate of 250 Hz and stored on a PC hard disk for offline processing and analysis. In parallel with the EEG data recording, two types of behavioral activity (active behavior/inactivity) were scored by an experienced observer. The animals were handled for a few seconds if a suspicion of sleep was observed (i.e., the animals did not move and tended to close their eyes).

### 2.2. Preprocessing

The first part of data preprocessing was done by an expert. Data were preprocessed in BrainVision Analyzer 2 software (Brain Products GmbH). At first, the data were manually inspected, and all artifacts were discarded. Then, the data were segmented by markers indicating behavioral (in)activity, and only epochs corresponding to behavioral inactivity were extracted as a model of resting state EEG data. Segments shorter than 2 s were excluded from further analysis. Data was then exported to be analyzed in Matlab. The second part of the data preprocessing was done by the EEGlab toolbox [20]. Here, the EEG recordings were re-referenced to average and filtered by using a 1-40 Hz two-way band-pass FIR filter with 2000 filter coefficients.

### 2.3. Microstates analysis

The microstate analysis was performed on 2-minute segments of broadband (1-40 Hz) EEG. Microstates were computed at all peaks of the global field power (GFP) [1,21] by the agglomerate hierarchical clustering algorithm (AAHC algorithm) [22, 23]. The microstate analysis was executed using the EEGLAB plugin for microstates, version 1.2, developed by Thomas Koenig [24]. The analysis was performed for different microstates ranging from 2 to 10. The global average topographies were used for backfitting to individual EEG recordings.

All typical microstate parameters (coverage, occurrence, duration, and GFP) were computed for microstate description. The strength of the average global activation during a given microstate *k* was defined by its average GFP of all EEG samples assigned to microstate *k*:

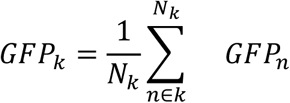

where *N_k_* is the number of samples assigned to the cluster *k* [25].

The duration of each microstate was represented by a time epoch, in milliseconds, during which all successive, original maps were assigned to the same microstate class, starting (ending) at the midpoint in time between the last original map of the preceding microstate and the first original map of the following microstate. The number of appearances of the microstates of a given class per second across all analysis epochs defined occurrence. Coverage was referred to as the meantime in milliseconds, which was covered across all analysis epochs by all microstates of a given class; this value can also be read as a percentage of the entire analysis time. [26]

The transition probabilities between the microstate classes were extracted as the asymptotic behavior of transitions between microstates (i.e., the likelihood of switching between different microstates) [27]. Expected transitions have been computed and are defined by the following equation:

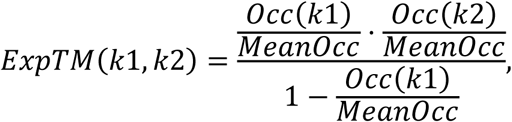

where *k*1, *k*2 represent the microstate number, *Occ* represents the occurrence, and *MeanOcc* represents the mean occurrence [24].

The above-mentioned parameters were proved to be effective markers for distinguishing different brain states in the case of human EEG [28].

### 2.4. The optimal number of microstate classes

For estimation of the optimal number of MS, several indexes were utilized based on the human studies: i.e Krzanowski-Lai (KL) criterion [22,29], the cross-validation criterion [4], or the metacriterion [7], which consists of several criteria. A list and description of the parameters that optimize the number of microstates can be seen in the supplements part, see Appendix 6.

The global explain variance (GEV), the percentage of data variance explained by a given set of microstate maps [22, 30]) was computed. The GEV value for a specific microstate map with label *k* is defined by the following equation [31]:

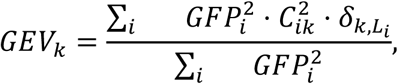

where *GFP_i_* is the global field power (defined as the standard deviation of the instantaneous EEG topography), 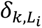 is the Kronecker delta, i.e., 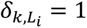 for *L_i_* = *k* and 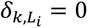 otherwise, and 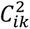 is the squared covariance between the instantaneous EEG topography and candidate microstate map. The total global explained variance (totGEV) is the sum of the GEV values over all microstate maps [31]:

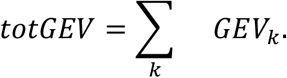

### 2.5. Generation of shuffled data

The surrogate data were generated to facilitate statistical permutation testing. The process involved generating a random vector of time positions for shuffling, repeated 10 times for each EEG record. Subsequently, the beginnings and ends of each electrode time series were swapped, resulting in temporal shuffling of the data. This method is based on the study by Mishra et al. [14], where a similar technique is used for the validation. The surrogates were then used for the *GEV* computation, and the statistical analysis of GEV values was performed to identify if there was a difference between the microstate models of surrogates and EEG data. If the microstate model captures EEG brain states specifically, there should be significantly higher GEV observed on the brain data compared to surrogates.

### 2.6. Spatial similarity of microstates

To quantify the spatial similarity between the identified microstate topographies, the Pearson correlation coefficient [32] was calculated. This was done to ensure that the polarity of topographies was ignored therefore the absolute value of Pearson’s correlation was taken.

### 2.7. Source localization of microstates

Following human EEG microstates studies [17,18], we aimed to determine the brain sources associated with the temporal evolution of individual microstates, considered as a representation of brain EEG activity. For that purpose, we generally followed a source localization pipeline introduced in our previous study [33]. The only difference from the previous study is that the 19-electrode cortical EEG system (instead of 12) was utilized to define the forward model. To estimate the sources, we applied the eLORETA inverse algorithm [34], implemented in the FieldTrip toolbox [35], to whole 2-minute-long segments extracted from each EEG recording as described in subsection 2.2. At first, three time series corresponding to three orthogonal directions were estimated for each source position. Second, three time series were projected onto the first principal component, which was further considered an EEG source time series for a given source position. To study the frequency specificity of underlying brain sources as in [17], each EEG source time series was further band pass filtered (two-pass Butterworth filter, order 3) into typical human EEG frequency bands: Delta (δ, 2 ≤ x <4 Hz), Theta (*θ*, 4 ≤ x < 8 Hz), Alpha (*α*, 8 ≤ x < 12 Hz) and Beta low and mid-range (*β*, 12 ≤ x < 20 Hz). Subsequently, the signal envelope (*BLP*) was computed by applying the Hilbert transform to each time series and taking the absolute value. The rationale behind associating microstates’ time series with frequency-specific signal envelopes is two-fold. First, the literature supports the idea that the underlying sources of human EEG microstates are frequency-specific [17]. Second, the signal envelope of a broadband signal is naturally driven by oscillations with the highest power, i.e., the δ frequency band, which may result in overlooking other frequency-specific information when associating with the microstates’ time series. To determine the brain sources associated with each of the five microstate topographies, we utilized and adapted a regression approach from [18]. For each subject *s* and the microstate *k*, a regressor *B_s,k_* was derived as:

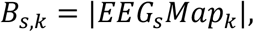

where *EEG_s_* is the EEG data matrix with dimensions (time samples x number of electrodes), and *Map_k_* is a single-session map of a microstate *k*. Note that for each frequency-specific signal envelope, the microstate regressor was the same. Deriving a microstate regressor this way is in contrast with [18] where the authors considered *a winner-take-all* property of microstates. Nevertheless, based on the work of [36] and rather slow spectral properties of the EEG signal envelope, it’s reasonable to derive a microstate regressor this way [37]. Furthermore, to eliminate the influence of global power signal fluctuation, we included two other regressors into a linear model: GFP and Global map dissimilarity (GMD) as in [18]. Therefore, for each source *s*, microstate *k*, and frequency band *f*, we estimated the parameters of the following linear model:

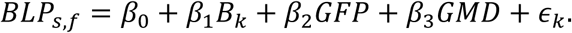

Finally, for each microstate *k* and frequency band *f*, we performed a group-level statistical analysis on *β*_1_ parameter to determine where in the brain and in which frequency band *B_k_* is related to the source-space BLP. The Monte Carlo permutation statistical analysis with the Threshold-free cluster enhancement (TFCE) method to correct for multiple comparisons [38] was performed utilizing the FieldTrip toolbox [35] to determine where *β*_1_ differs from zero (α = 0.001, two-sided test). To associate clusters where the difference of *β*_1_ from zero is mostly pronounced to rat brain areas, we also coregistered an anatomical rat brain atlas from [39] to label each source-space position to a corresponding rat brain anatomical area. Obtained statistical maps were subjected to two summary statistics. First, we computed the total cluster size by simply dividing all sources surviving TFCE correction by the total number of sources, which represents spatial as well as frequency specificity of the given microstate association with source-space BLP. Second, a correlation matrix of each frequency-specific microstate statistical map (TFCE correction masked) was computed to evaluate intra-as well as inter-microstate frequency-dependent statistical source map similarities.

## 3. Results

### 3.1. The optimal number of microstates

The results of estimating the optimal number of microstates utilizing CV, KL, Davien Bouldin, Dunn, Frey, and Van Groenewould, dispersion, and metacriterion methods showed substantial differences in estimating the number of MS classes. As the KL method exhibits insensitivity to the number of electrodes [40] and is frequently used in human studies [41–48], the outcome of KL was set as the preferred choice for the current study, resulting in an optimal number of microstates established at five. More details on the assessment of MS classes are provided in Supplementary material, see Appendix A and Figure 8.

### 3.2. Validation on shuffled data

The GEV parameter was computed for both the original broadband and shuffled data. The outliers were excluded from both groups. An outlier is a value of more than three scaled median absolute deviations (MAD) from the median. Boxplots of mean GEV and histograms for each number of microstates (from 2 to 10) are plotted in (Figure 2B).

**Fig. 2.**
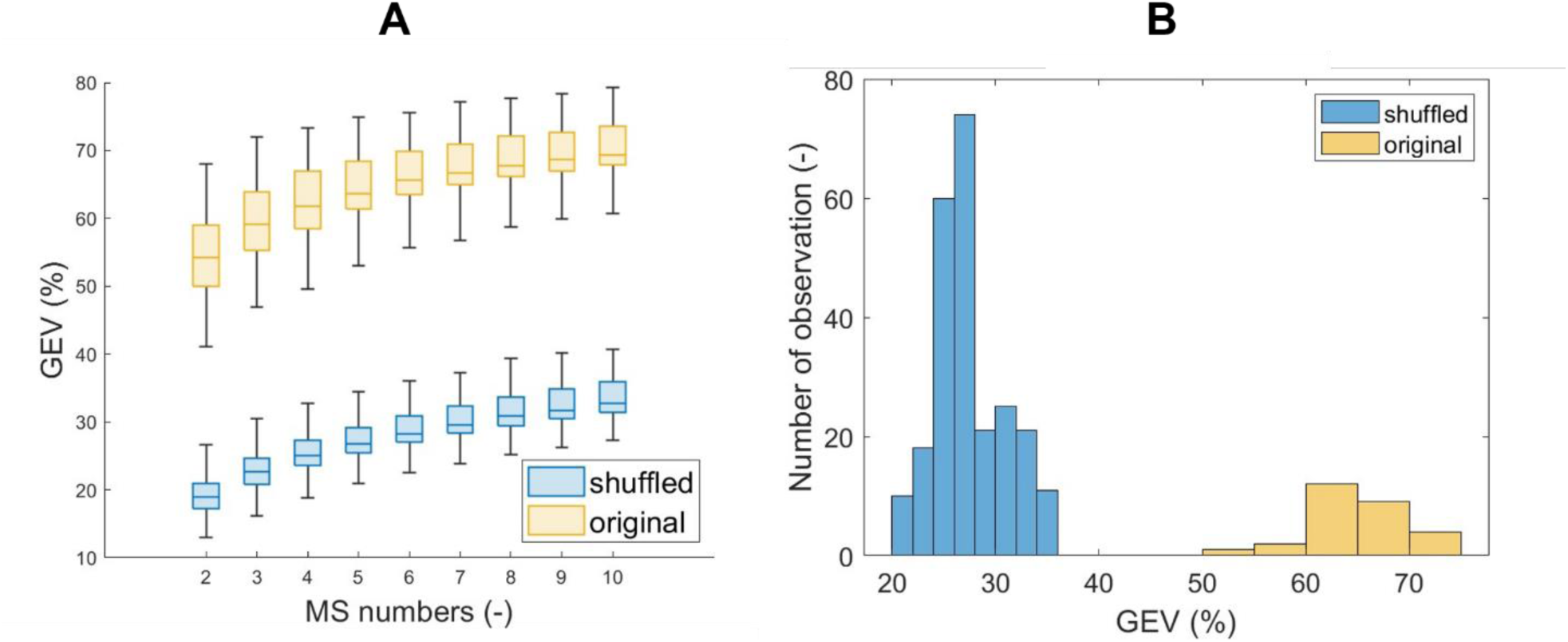
The GEV values for the original dataset are depicted in yellow, while those for the temporally shuffled dataset are in blue across various numbers of microstates. Panel A illustrates boxplot graphs for microstates 2 to 10. Panel B shows the histogram of GEV values for the optimal number of microstates set at 5.

Normality testing using the Kolmogorov-Smirnov test revealed that neither the whole shuffled dataset nor the shuffled dataset for five optimal microstates followed a normal distribution (p-value < 0.001). The same non-normal distribution pattern was observed for the original whole dataset and the original dataset for five optimal microstates (p-value < 0.001). The two-sample Kolmogorov-Smirnov test confirmed (at an alpha level of 0.05) that the data did not stem from the same distribution (p-value < 0.001). Statistically significant differences between the distributions were identified for both the entire datasets and five optimal microstates.

GEV values for the original broadband dataset were, on average, 64.81±7.06 (mean ± STD) for microstates numbers from 2 to 10 and 64,72 %±4,94 % for 5 microstates specifically. In contrast, for the temporally shuffled dataset, average GEV values were 30.10±7.31 (mean ± STD) for microstates from 2 to 10 and 29.55±6.00 for five microstates correspondingly. The significant decrease in GEV values after temporal shuffling suggests that the outcomes of microstate analysis are non-random.

### 3.3. Microstate description

The topographical representation of five extracted MS classes and their spatial correlation coefficients are presented in Figure 3B. MS1 and MS3 display similar anterior-posterior topography; MS2 is centered on the midbrain and further spreads to the parietal posterior regions. MS4 is represented by a both-sided temporal activation connected on a mid-line, and MS5 is represented by a central locus. Topographies and spatial correlation plots for the number of microstates from 2 to 10 are presented in Supplementary material for information purposes, see Appendix.

**Fig. 3.**
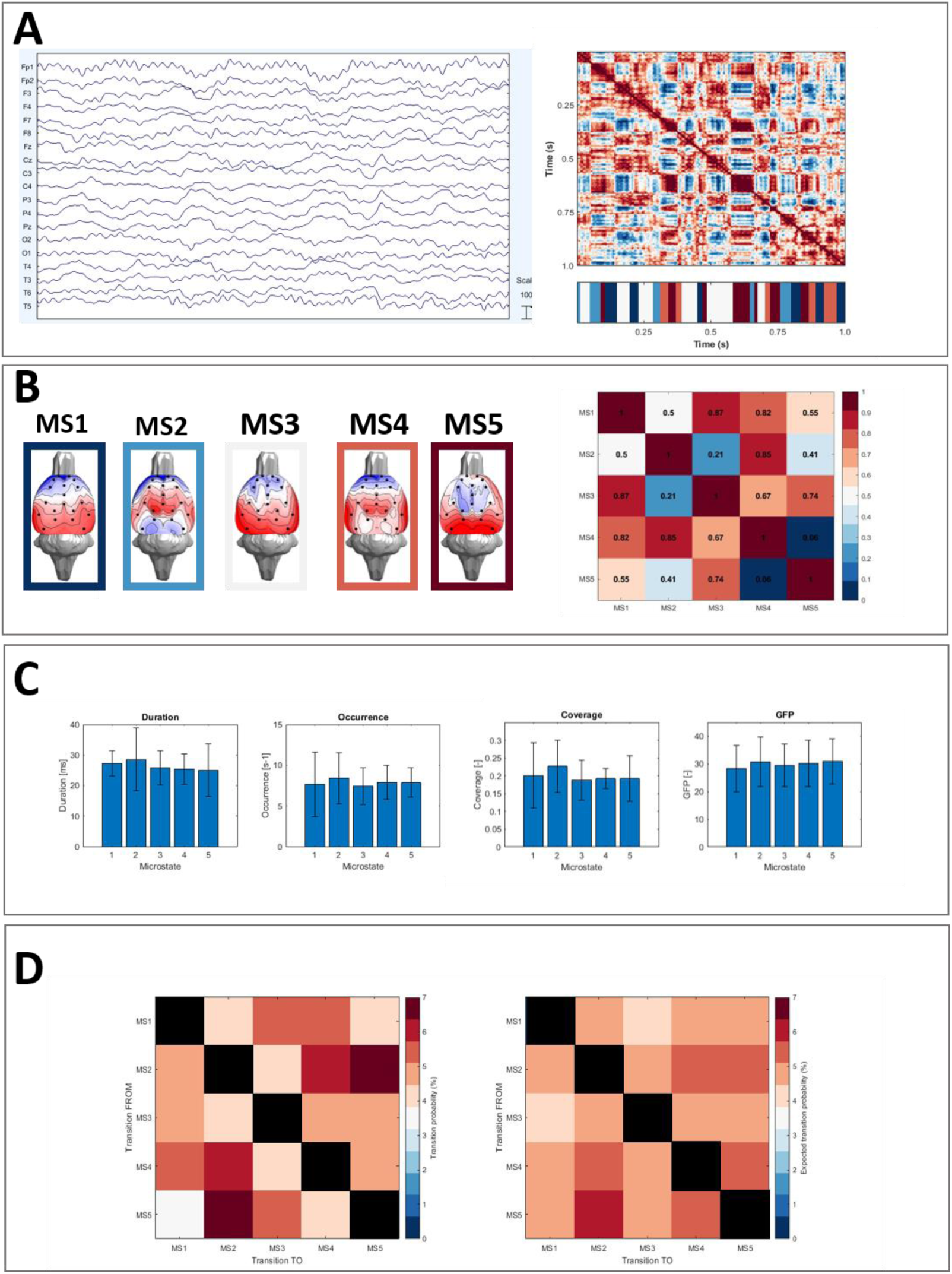
The results of microstate analysis on rats’ EEG. Panel A represents the preprocessed rats’ EEG and the spatial correlation of the topographic maps between time samples. Panel B depicted the resulting five microstate topologies and the spatial correlation between the microstate topographies. Panel C describes the classical microstate parameters, concretely their mean and standard deviation values for each microstate. Panel D transition probabilities between microstates: the left matrix shows observed probabilities (in percentage) and the right matrix illustrates expected transitions based on occurrences.

The details of classical parameters of MS analysis for five microstates are presented in Figure 3C. As can be seen, the average coverage of microstates was at around 0.20, and the duration was measured at around 26 ms. Transition probabilities between extracted microstates are plotted in Figure 3D.

### 3.4. Source Localization of Microstates

We computed group-level statistical maps representing the association between each microstate time course and frequency-specific source-space BLP signal fluctuations. Figure 4A visualizes a percentage of the total cluster size from the total amount of sources for each microstate and frequency band. Figure 4B shows pair-wise correlation coefficients of each frequency-specific microstate map. Rows and columns corresponding to statistical maps not having significant activations are omitted.

**Fig. 4.**
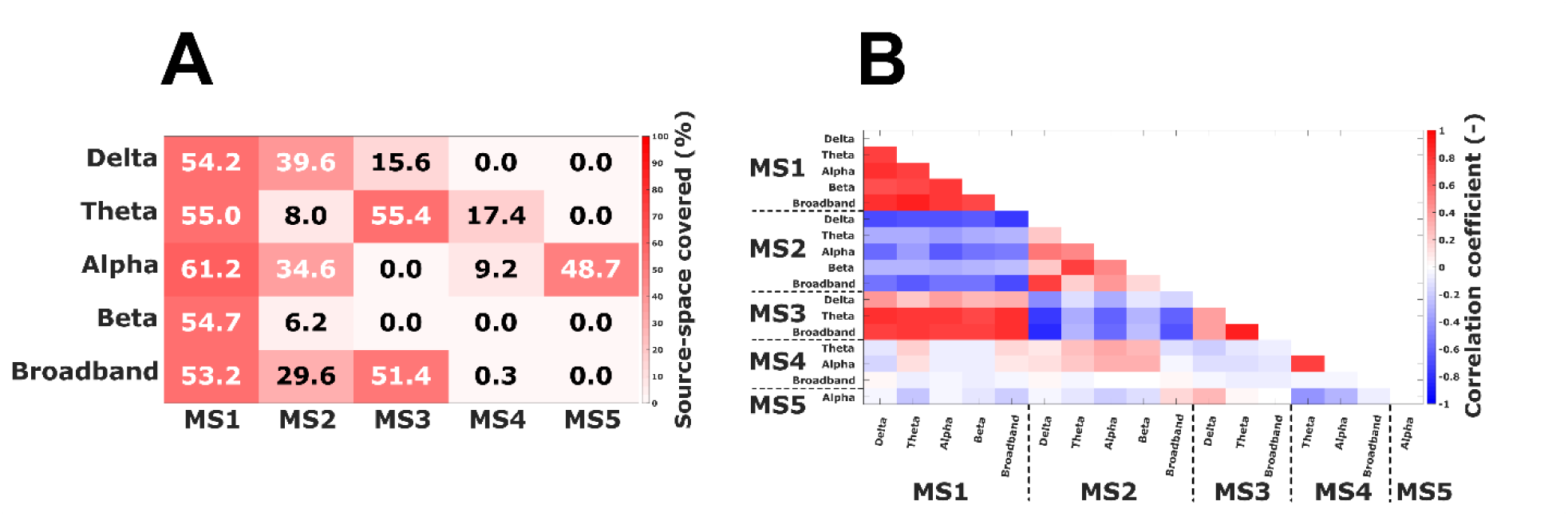
Source localization summary statistics for: **(A)** A activation level of microstates in the brain (as a percentage of sources included within clusters to the total number of sources) and **(B)** A Pearson correlation analysis of frequency-dependent source maps (both within and between microstates). Note that rows and columns corresponding to non-significant maps are omitted.

In Figure 5, the actual source maps are visualized. We can observe that MS1 is not frequency-specific, but its time series is rather associated with source activity in each frequency band. Spatially, clusters cover more than half of the source space, and the statistical maps seem to correlate strongly between frequency bands. MS1 extends over the entire thalamus, basal ganglia, and insular cortex over all frequency bands. In the case of MS2, the association with source activity is more pronounced in delta and alpha frequency bands as well as broadband. The broadband statistical map is more similar to the delta statistical map than to other frequency-specific maps. All MS2 statistical maps are negatively correlated with all MS1 statistical maps. In the delta band, or rather, in the broadband, MS2 occupies a similar space as MS1 but is localized slightly lower, extending to the ventral pallidum and hypothalamus. The theta to the beta band shows the greatest activity frontally - prelimbic cortex and frontal association cortex, and then in the primary and secondary auditory cortex. Interestingly, MS3 is widely associated with the source-space activity in theta and broad-band frequency bands, which are also strongly intercorrelated. MS3 patterns are generally anti-correlated with all statistical maps of MS2 and positively associated with MS1 statistical maps, as it replicates MS1 in its sources; see Figure 7. In the delta band, however, MS3 defines a narrow band of granular insular cortex, retrosplenial dysgranular cortex, and periaqueductal gray activity. MS4 expresses alpha and theta activity in frontal association, insular and infralimbic cortex, and its statistical maps are highly correlated. Conversely, MS4 maps are less similar to other MSs statistical maps. Finally, MS5 has only an alpha band-specific source-space pattern that is, again, less similar to patterns of other MSs compared to MSs 1-3. It extends over large areas of the primary and secondary auditory, entorhinal, dysgranular, and granular insular cortex.

**Fig. 5.**
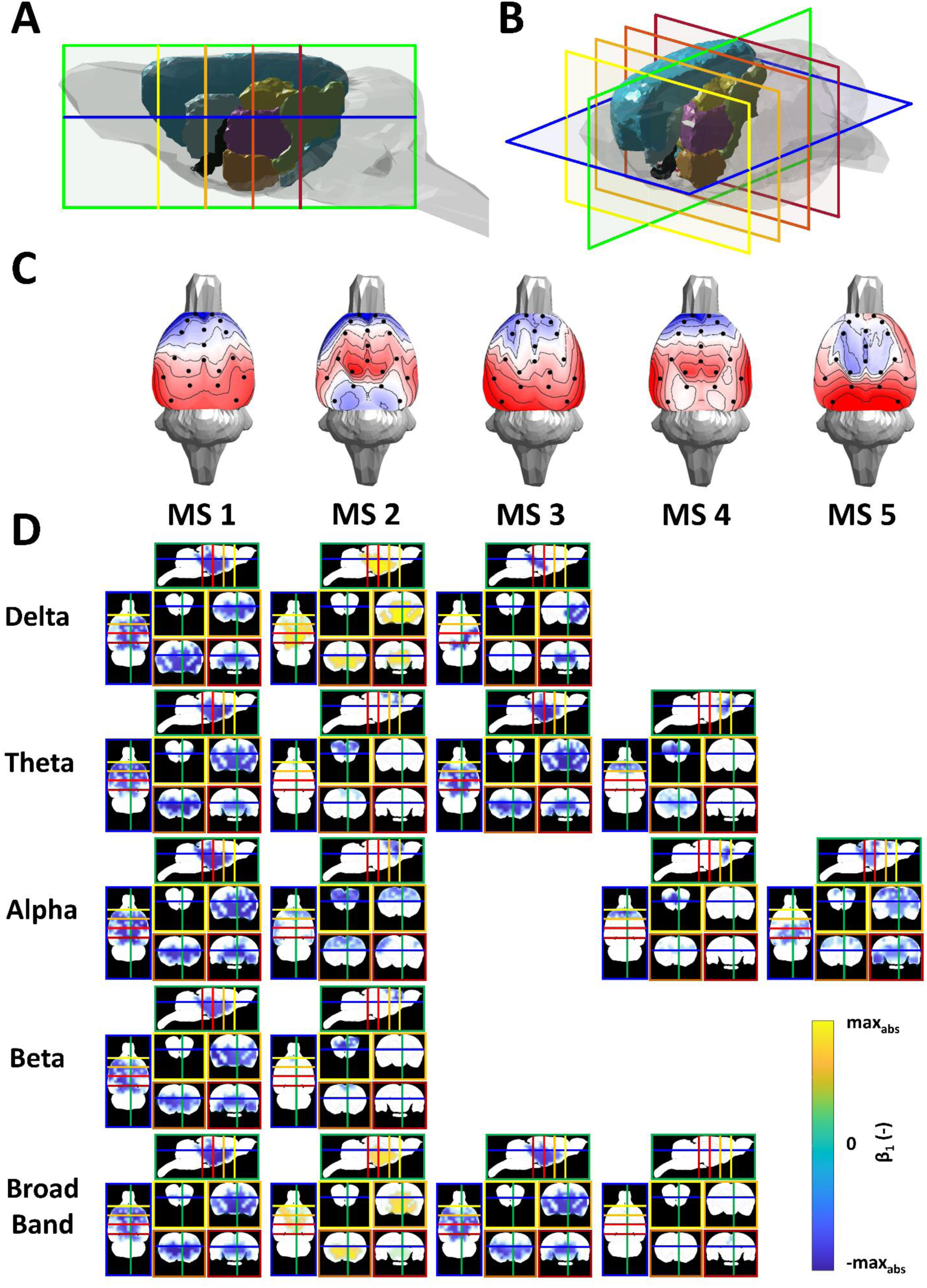
Sources of microstates. **(A,B)** A 3D model of the rat brain showing selected cutting planes for source presentations from right **(A)** and top right **(B)** view. The planes are: horizontal (−6 mm ventro-dorsally, framed in blue), sagittal (+0.5 mm laterally, framed in green), and four coronal sections (+3.5 mm anteroposteriorly, framed in yellow; 0 mm, in light orange; −3.5 mm, in dark orange; −7 mm, in red), coordinates refer to bregma according to [19]. Structures shown: Isocortex (Light Blue), Pallidum (black), Hippocampus (golden yellow), Hypothalamus (coral red), Diencephalon (orchid pink), Midbrain (light green), Striatum (light cyan) **(C)** Topographic maps of five microstates. **(D)** Group cluster permutation statistic values (ꞵ_1_, p < 0.001) interpolated on rat brain anatomical MRI scan shown in the respective planes with microstates’ classes in columns and the frequency bands in rows.

## 4. Discussion

In this study, we show that resting-state EEG from freely moving rats can be decomposed into multiple microstates resembling those described in humans. By applying various criteria to determine the optimal number of microstates and choosing the Krzanovski-Lai (KL) method, we identified five microstate classes. Validation against shuffled data confirmed that these microstates represent non-random temporal patterns, as evidenced by significantly higher Global Explained Variance (GEV) in the original dataset compared to shuffled data. Each of the five microstates displayed distinct scalp topographies and classical parameters (duration, occurrence, coverage, and Global Field Power), with transition probabilities suggesting certain preferred transitions. Source localization revealed specific frequency-dependent activation patterns for each microstate, highlighting both cortical and subcortical regions and underscoring the translational potential of microstate analysis for understanding large-scale brain dynamics in animal models.

In human studies, four maps have traditionally been recognized [8, 30]; however, the development and implementation of more sensitive criteria resulted in a recommendation not to strictly adhere to the number of four [18]. Recent studies have demonstrated that the optimal number of microstates describing resting-state EEG varies from 4 to 7. Using the Krzanovski-Lai criterion that is insensitive to the number of electrodes [40] and frequently used in human studies [49], we identified five microstate maps explaining 71% of the variance, somewhat less than the variability explained in human studies, which is commonly around 80% [3]. Although the actual values of explained variance are not directly comparable between rats and humans, one possible explanation for observing lower explained variance in rats is that cortical EEG recordings are unaffected by the blurring effects of skull and scalp tissues. This results in a richer signal that may be less effectively captured by the same number of microstates as in scalp EEG studies. It is also worth noting that most human research is conducted with participants’ eyes closed. Studies investigating EEG microstates during eyes-open conditions have shown that, while the topography and relative proportions of parameters remain largely unchanged, there is typically a decrease in overall measures such as GEV, duration, and time coverage [52, 64]. Consequently, the fact that our rats’ eyes were open may have contributed to the observed findings.

In human classical 4-map arrangement, microstate map A exhibits a left-right orientation, map B a right-left orientation, map C an anterior-posterior orientation, and map D shows a fronto-central maximum. Even if more cluster maps are selected, these four canonical maps seem to consistently dominate the data across different age ranges, conditions (e.g., sleep and hypnosis), and pathological states [1]. Here, the estimated rat microstates were also characterized by distinct dominant topographies (Figure 3). The observed spatial correlation between topographic maps across the time samples further supports the idea of microstates in rats. In Figure 3c, clear temporal clusters of topographically similar EEG activity can be observed. MS1 exhibits a frontal to occipital topography and is highly correlated with MS3 (pronounced occipital activity) and MS4 (fronto-central and both-sided temporal activation); MS2 was characterized by central activation and demonstrated strong spatial correlation to MS4; finally, MS5 resembled a centro-parietal locus of activity. The identified microstates showed a high number of spatial inter-correlations, a finding commonly reported in human data [18]. Although none of the rat microstates show a lateralized form, as is true for human microstates A and B, based on spatial similarity, we might infer that rat microstate 1 is most similar to human microstate C [4, 7], in humans associated with control of cognitive processes [8, 50]. Rat microstate 5 could then represent human microstate D [4, 7], which in humans is associated with the vigilance level [50–52].

Among the four rodent studies capturing microstate-like activity, only Mishra et al. [14] reported results in a traditional way that allows for parameter comparison. However, they used microelectrode LFP recordings, targeting some kind of mesoscale microstates, while we employed a superficial electrode system targeting areas homologous to those studied in humans. Mishra et al. provided GEV values against surrogates −0.71 for the prefrontal cortex and striatum and 0.64 for the ventral tegmental area, which aligns closely with our findings. Although, like Mishra et al. [14], we observed a significant decrease in GEVs compared to the surrogate data, Jajcay et al. [63] showed, for example, that in the case of surrogates based on vector regression models and Fourier transforms, the microstates were not statistically distinguishable from them. Moreover, Pascual-Marqui et al. [64] showed that microstate properties can be effectively captured by a multivariate cross-spectrum EEG model. Thus, caution is advised when interpreting microstate findings in relation to surrogate data.

The mean temporal coverage values (0.23 for PFC, 0.23 for STR, and 0.26 for VTA) in Mishra et al. [14] are also consistent with our result of 0.20. However, the key difference emerged in the average duration of microstates: 26.99 ms in our study compared to 68.04 ms for PFC, 70.30 ms for STR, and 65.41 ms for VTA in Mishra et al., i.e. values that are more similar to those found in human research (60–120 ms). Despite this difference, our results had lower standard deviations, and our durations still fell within the predicted range for human microstates (20.7 to 106.8 ms) [49]. The shorter durations in our study likely led to a higher occurrence rate of microstates, approximately twice that observed by Mishra et al. Importantly, the choice of four microstates in their study may have contributed to differences in reported parameters, as variations in the number of microstates are known to influence both duration and occurrence rates [7]. As indicated by the transition probabilities between microstate classes, certain patterns of the activity were more stable than the others - for example, the bidirectional transitions from MS2 to MS5 and MS2 to MS4 were the most frequently occurring ones, showing that the dynamics between microstates are likely to be unique, as in humans [27, 52]. However, this aspect remains difficult to compare due to a lack of normative data on microstate transition probabilities in humans.

We estimated the neuronal generators of five resting-state-like topographies using a recently developed source-localization approach [33] and temporal general linear modeling inspired by the study of [18]. This analysis identified several areas of activation shared across three microstate classes (MS1-MS3), with each of the five microstates consistently linked to distinct frequency band-specific brain regions. The overlap in brain areas across microstate source spaces likely explains the partial spatial similarities in scalp potential maps (i.e., resting-state-like topographies).These shared regions include areas along the anterior and posterior medial axes, the insula, and the superior parietal cortex. Custo and colleagues also identified a set of brain regions active across most microstate networks in human microstates [18]. These regions, including the anterior and posterior cingulate cortices, precuneus, superior frontal cortex, supramarginal gyrus, dorsal superior prefrontal cortex, and insula, are key hubs frequently highlighted in studies of structural and functional brain networks [53–55].

Previous human studies have linked EEG microstates to fMRI BOLD signals to associate large-scale brain networks with specific microstates [8, 56]. Microstate A was connected to the auditory network, microstate B to the visual network, microstate C to the salience network, and microstate D to the attention network, with corresponding brain regions. While many studies have identified large-scale rat brain networks, recent debates on the overlap and subdivision of fMRI resting-state networks suggest caution when linking microstates to specific brain functions based solely on fMRI correlations [7]. In our study, MS1 showed the most prominent activity within the thalamus, basal ganglia, and insular cortex; MS2 shared similar locations; however, it was centered lower, also reaching ventral pallidum, hypothalamus, prelimbic, frontal association, and auditory cortices. MS3 also shared regions with MS1, including granular insular and retrosplenial cortices with periaqueductal gray. MS4 was specific for the frontal association, insular and infralimbic cortices, MS5 then for auditory, entorhinal, dysgranular, and granular insular cortices.

Milz and colleagues [17], unlike the others [8, 56], provided conclusive results on the interrelationship between EEG microstate classes and spectral power characteristics and showed that human microstates are driven predominantly by alpha activity. Our results show that rat head-surface topography of microstate classes appears to be determined by intra-cortical sources of specific frequency bands. While microstates 1 and 2 seem to be generated by activity within the broad frequency range, MS3 seems to be low frequency specific, MS4 is typical for the theta-alpha frequency range, and finally, microstate 5 is purely alpha band specific.

By employing the rat cortical EEG source localization technique [33], we can analyze the results at the level of specific brain regions, further enhancing the interpretability of the obtained microstate maps. However, it is crucial to recognize the inherent challenges associated with the EEG inverse problem [57]. Additionally, since the electrodes are primarily situated on the top part of the brain cortex, this setup limits the resolution for detecting activity in deeper brain structures. Consequently, the reduced accuracy in localizing signals from deep brain regions has to be considered.

## 5. Conclusion

The growing interest in non-invasive tools for evaluating brain function in various disorders highlights the need for translational methods applicable to both humans and animals. In this study, we aimed to identify and thoroughly describe microstates in the resting-state EEG of freely moving rats, which could be comparable to those observed in humans. Resting-state EEG microstates are a widely used and promising technique for assessing brain function in both healthy and pathological conditions in humans, but this approach has been largely overlooked in animal studies.

This study provides the first comprehensive identification and description of microstates in the resting-state EEG of freely moving rats, with findings that resemble human-like microstates. By using a classical microstate analysis pipeline and source localization techniques, we identified five microstate EEG maps in rats, explaining 71% of the variance in our dataset. These results align with human studies and suggest that microstates in rats might be linked to brain networks similar to those in humans, speculatively also to cognitive control and vigilance. The shared brain regions identified across microstates in our source localization analysis, including key hubs like the cingulate cortex, precuneus, and insula, further highlight the translational potential of this approach. Although care must be taken when linking microstates to specific brain functions, this study opens new avenues for cross-species research on brain function using microstate analysis.

## Acknowledgment

This study was performed and enforced based on the diploma thesis of Petra Peskova from the Faculty of Biomedical Engineering, Czech Technical University in Prague, with the topic: Evaluation of animal EEG recordings by microstate analysis.

## Author contributions

VP: conception and design of the work, microstate analysis, data interpretation; ČV: acquisition, data interpretation, and drafting the work; PP: analysis; MP: analysis, and drafting the work; SJ: source localization analysis, and drafting the work; VK: analysis; IGB: conception and design of the work, data interpretation, and drafting the work; TP: conception and design of the work, drafting the work, revising it critically for important intellectual content, final approval of the version to be published, agreement to be accountable for all aspects of the work in ensuring that questions related to the accuracy or integrity of any part of the work are appropriately investigated and resolved.

