## Supplementary Material for "Microstate in rat’s EEG: a proof of concept study"

### Appendix

#### Appendix A Description of the criterion

The meta-criterion combines several criteria, each operating on a slightly different principle of determining the optimal number of microstates. Based on all criterion estimates, the interquartile range (IQR) and the interquartile mean (IQM) are calculated [18]. The meta-criterion can include for example, followed criteria: Davies and Bouldin [58], KL [29], Dunn's [59, 60], Frey and Van Groenewoud's criterion [60, 61], dispersion [25].

The most commonly used one is the cross-validation criterion (CV), which was introduced in Pascual-Marqui et al. [21]. This measure is related to residual noise, and the goal is to obtain a low value of CV, which is defined by the following equation:

$$CV = \delta^2 \cdot \left(\frac{C-1}{C-K-1}\right)^2,$$

where $\delta^2$ defines the estimation of variance of the residual noise, $C$ is the number of the EEG channels and $K$ is the number of clusters [21, 25]. However, because the CV is a ratio between GEV and the degrees of freedom for a given set of template maps, this criterion is highly sensitive to the number of electrodes in the montage [22].

The KL criterion was introduced in Krzanowski and Lai 1988 [29], as a means of selecting how many clusters to use based on the dispersion measure. High values of KL usually indicate an optimal number of clusters [25]. It works by first computing a quality measure of segmentation, termed dispersion ($W$), which is defined by equations:

$$W_k = \sum_{k}^{K} \frac{S_k}{2 \cdot N_k},$$

where

$$S_k = \sum_{n}^{N} \sum_{n\prime}^{N\prime} \parallel x_n - x_{n\prime} \parallel^2,$$

for $l_n = k \wedge l_{n\prime} = k$ . Variable $S_k$ represents the sum of pair-wise distance between all maps of given cluster $k$ and $N_k$ is the number of maps for cluster $k$. The following equations then compute the KL criterion:

$$KL(K) = \left|\frac{DIFF(K)}{DIFF(K+1)}\right|,$$

$$DIFF(K) = (K-1)^{\frac{2}{C}} W_{K-1} - K^{\frac{2}{C}} W_K,$$

where $W_k$ represents the dispersion as is described in previews equations. The $KL$ criterion attains large values when an elbow in the $W_k$ curve occurs. [22, 25]

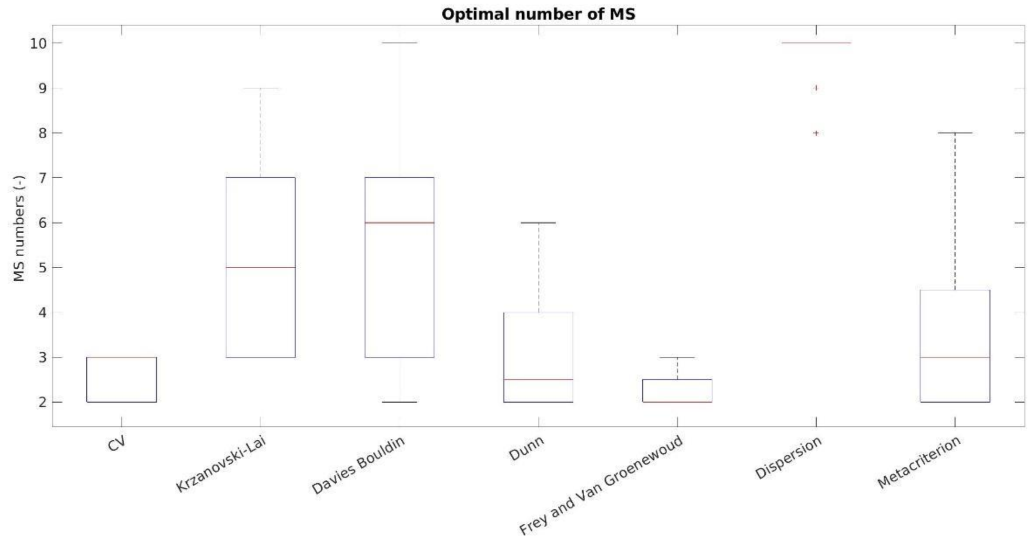


**Fig. A1** *Optimal number of microstates based on several criteria.*

#### Appendix B Changes in CV and GEV

The CV and GEV parameters were computed for the total number of microstates from 2 to 10. The mean value and standard deviation have been computed for both parameters and are depicted in Figure 9.

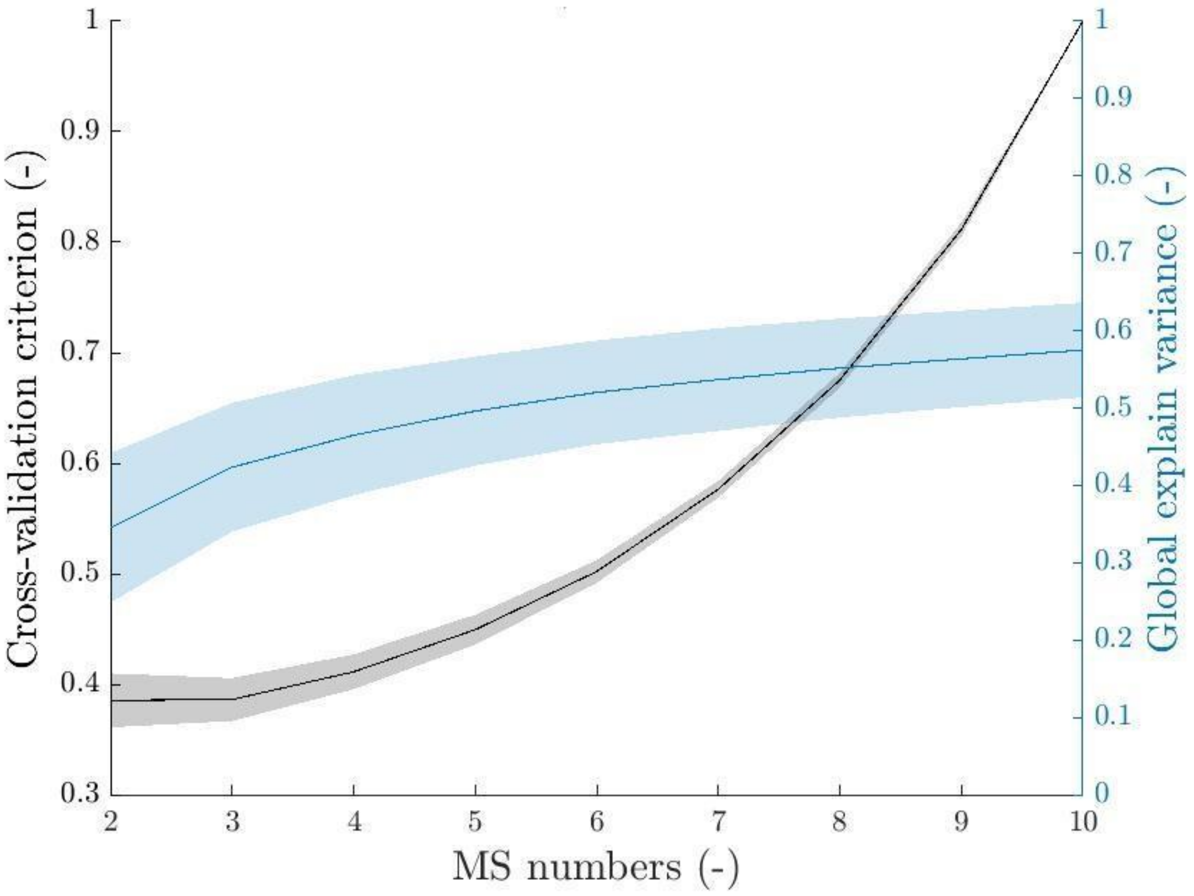


**Fig. B1** *The cross-validation (in black) and GEV (in blue) parameters across subjects are depicted here. The mean and standard deviation are presented for each variable.*

### Appendix C Microstate topography across the different MS number

This appendix reports the microcrastes' topographies for different numbers of microstates and their spatial correlations between them. Microstate analysis was performed as described in section 2.

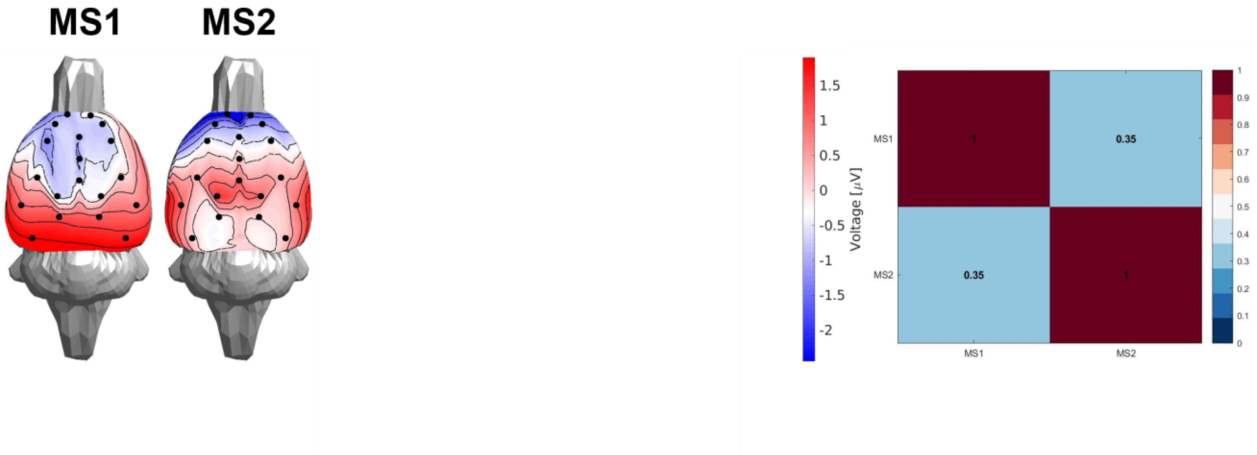


**Fig. C1** *The topography of microstate in case of two microstate total. The right panel defines the spatial correlation between the microstates.*

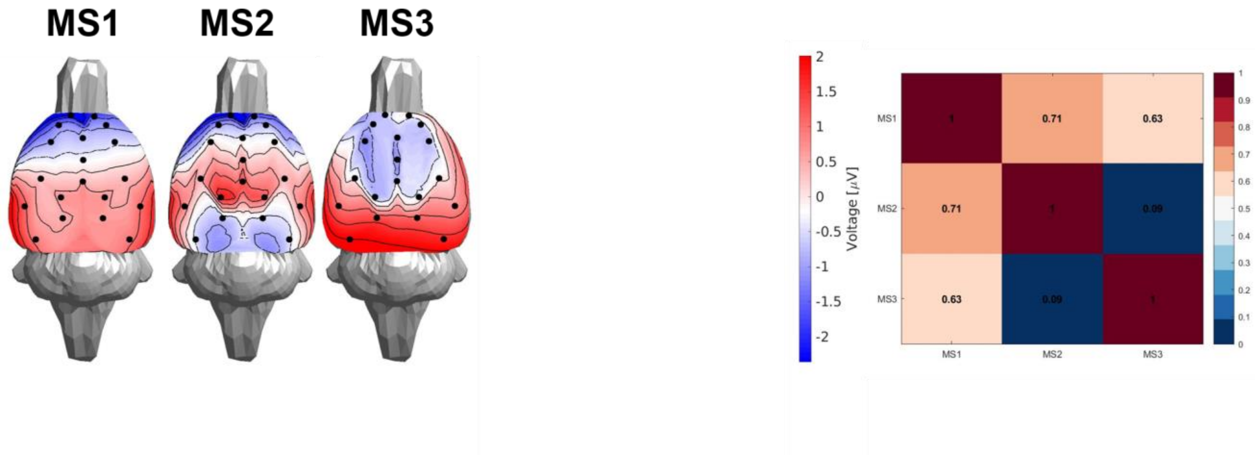


**Fig. C2** *The topography of microstate in case of three microstate total. The right panel defines the spatial correlation between the microstates.*

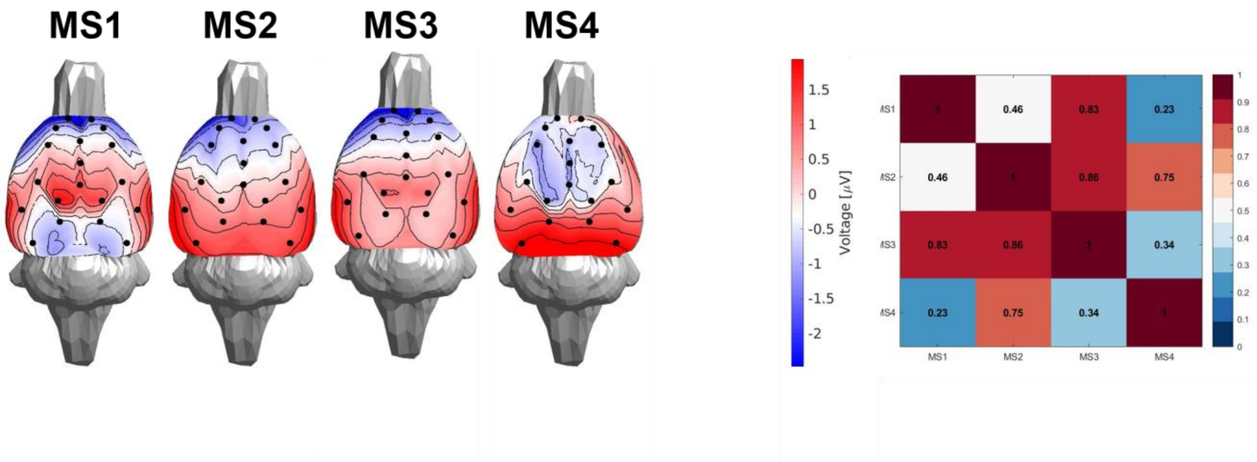


**Fig. C3** *The topography of microstate in case of four microstate total. The right panel defines the spatial correlation between the microstates.*

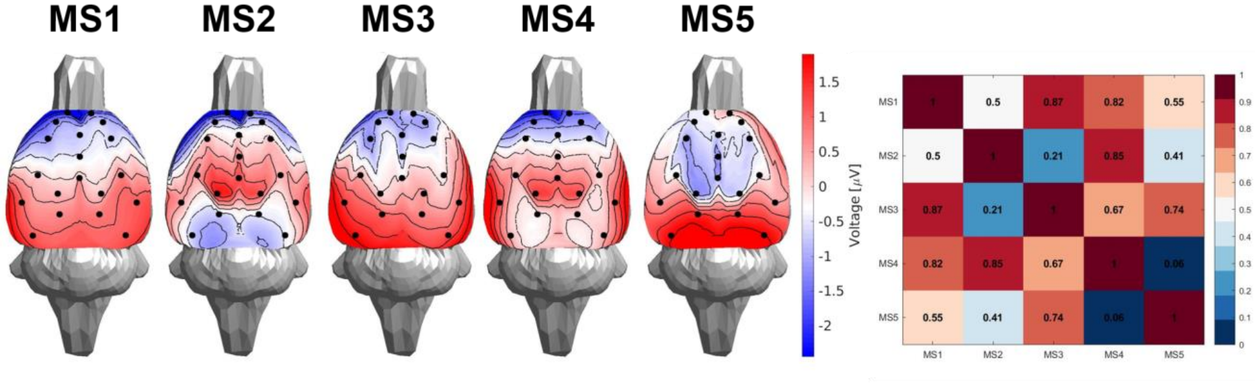


**Fig. C4** *The topography of microstate in case of five microstate total. The right panel defines the spatial correlation between the microstates.*

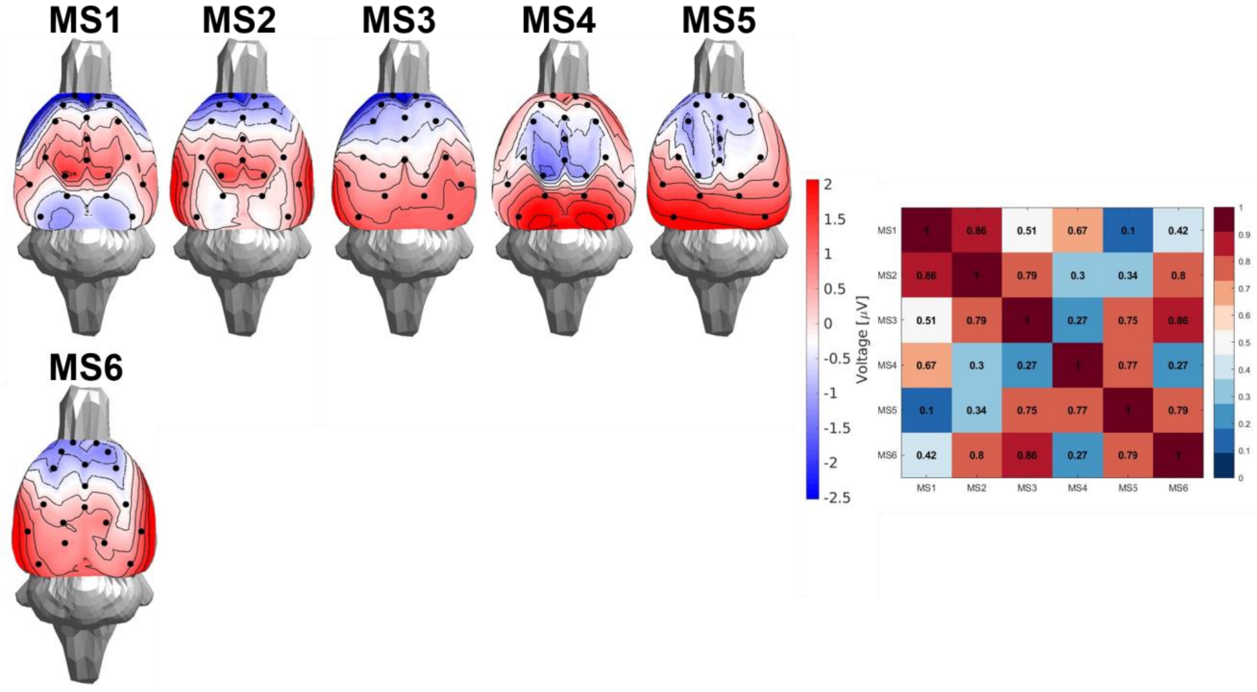


**Fig. C5** *The topography of microstate in case of six microstate total. The right panel defines the spatial correlation between the microstates.*

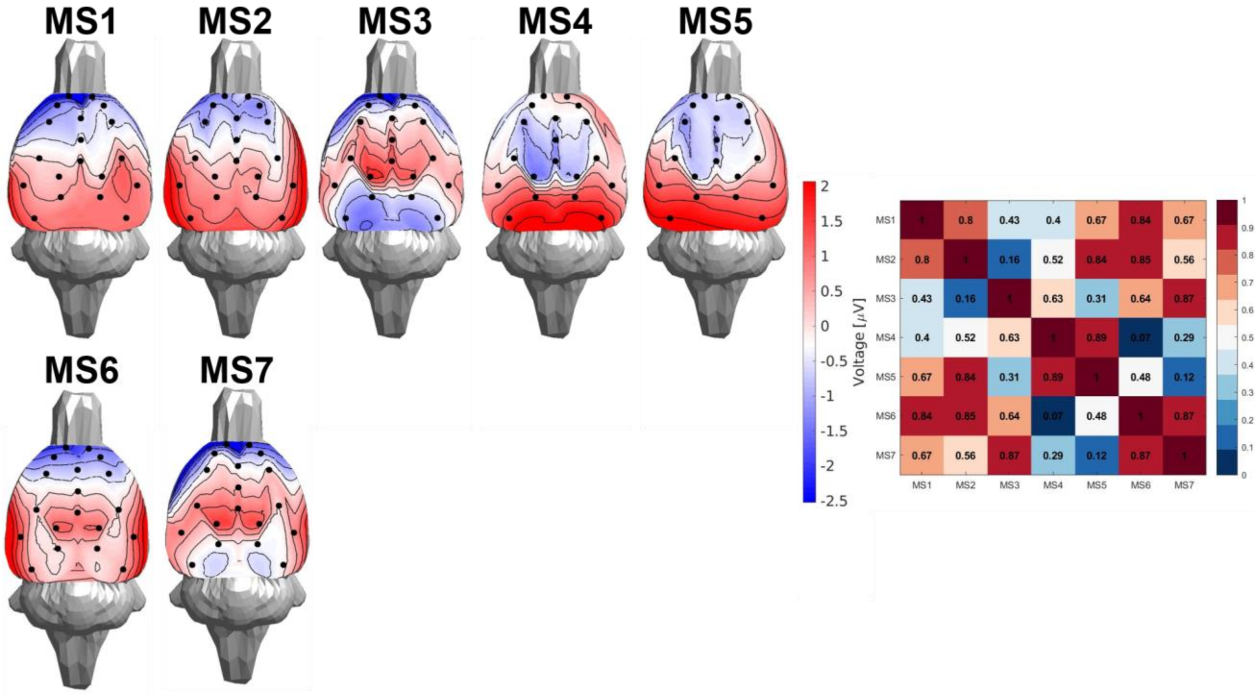


**Fig. C6** *The topography of microstate in case of seven microstate total. The right panel defines the spatial correlation between the microstates.*

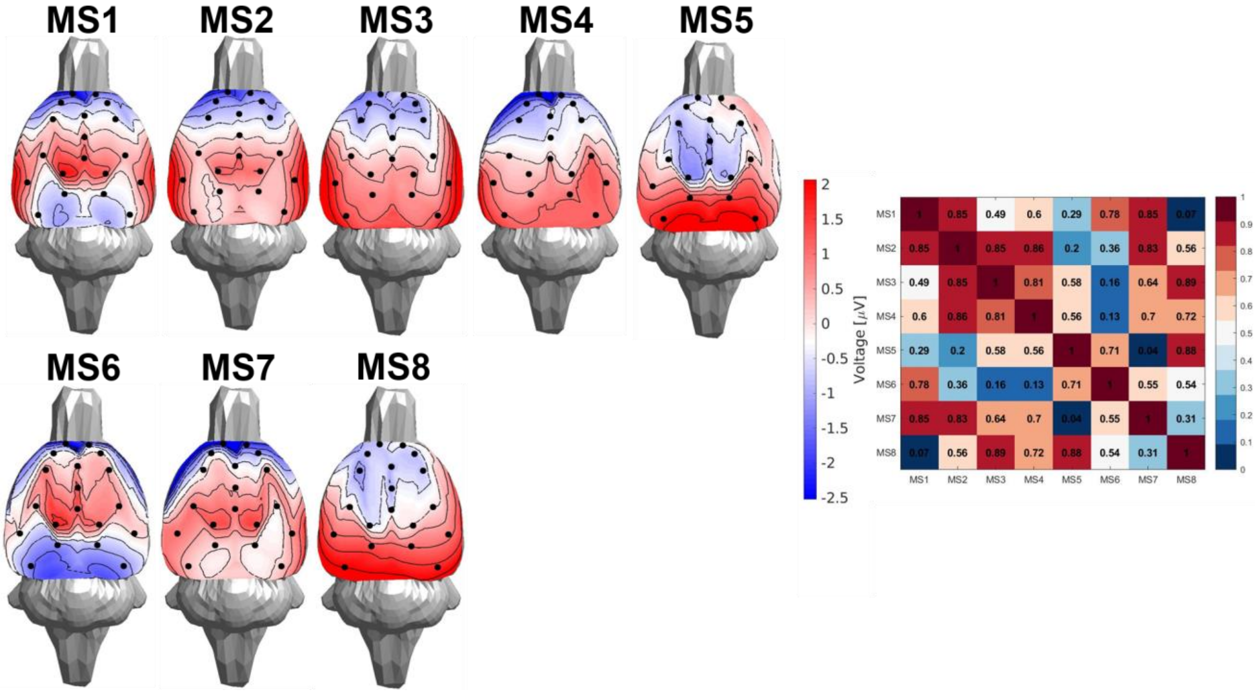


**Fig. C7** *The topography of microstate in case of eight microstate total. The right panel defines the spatial correlation between the microstates.*

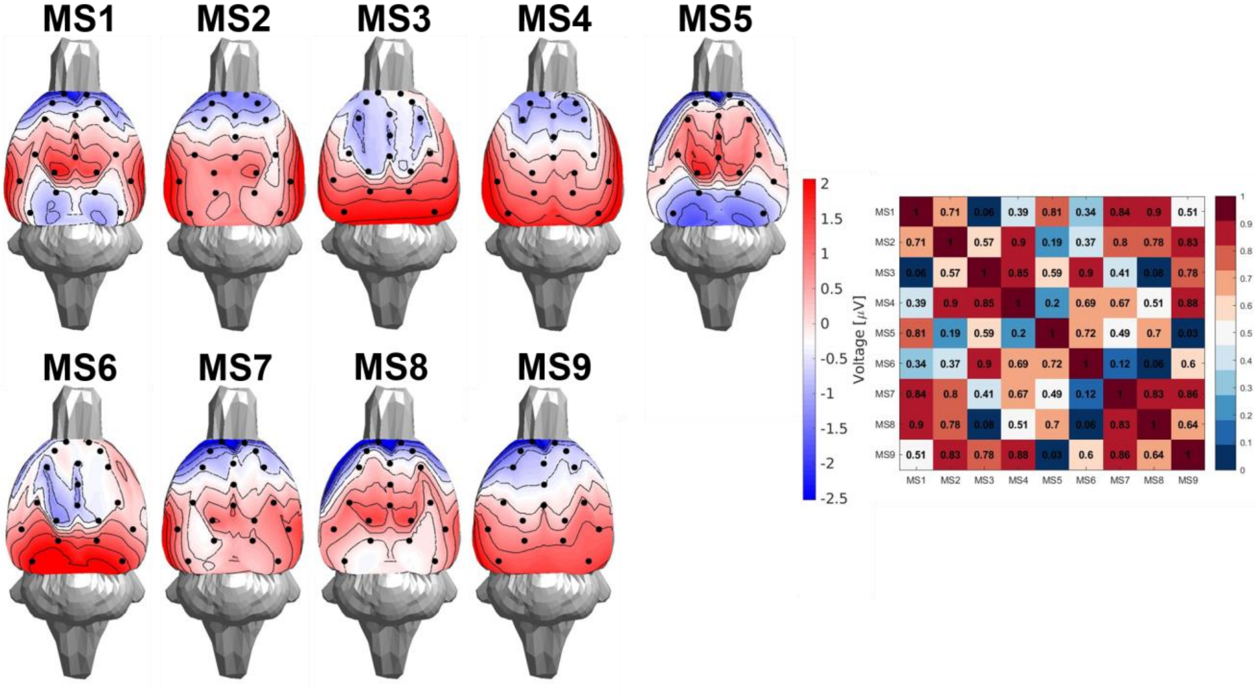


**Fig. C8** *The topography of microstate in case of nine microstate total. The right panel defines the spatial correlation between the microstates.*

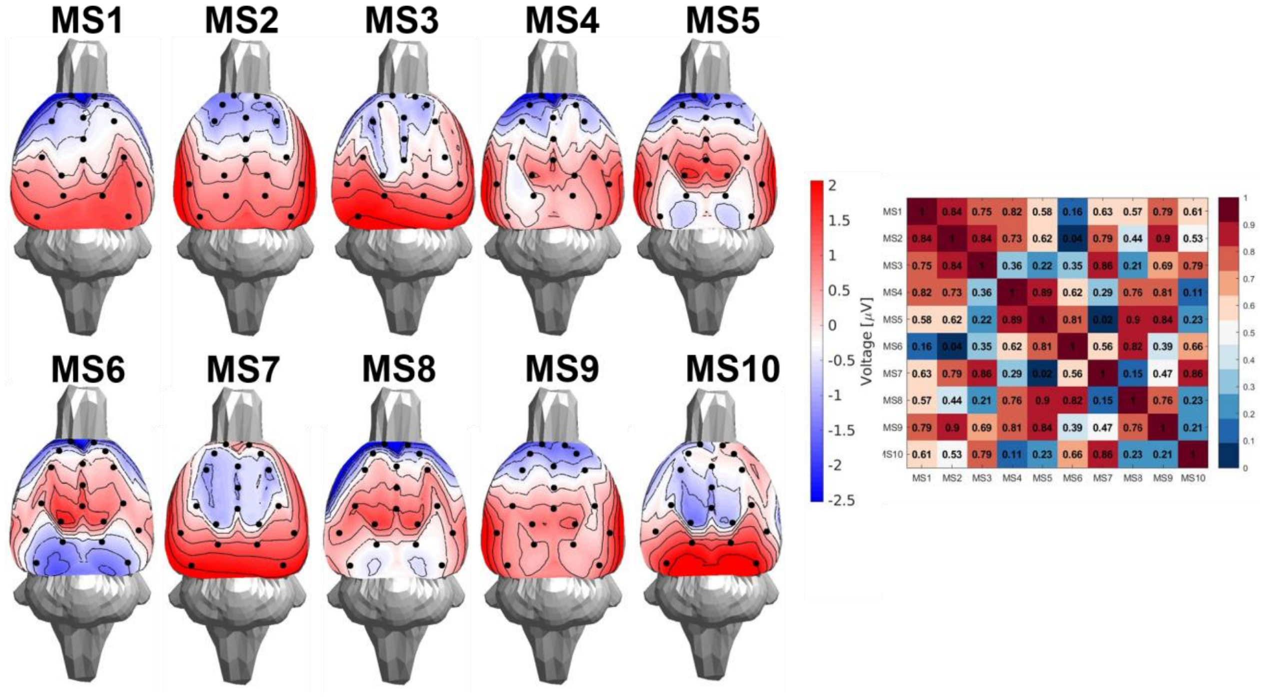


**Fig. C9** *The topography of microstate in case of ten microstate total. The right panel defines the spatial correlation between the microstates.*
